## Supplementary Information for "Breaking antimicrobial resistance by disrupting extracytoplasmic protein folding"

**This file includes:**

Legends for Supplementary Tables 1 and 2
Supplementary Tables 3 to 7
Supplementary references

**Supplementary Table 1. Analysis of the cysteine content and phylogeny of all identified β-lactamases.** 6,649 unique β-lactamase protein sequences were clustered with a 90% identity threshold and the centroid of each cluster was used as a phylogenetic cluster identified for each sequence (“Phylogenetic cluster” column). All sequences were searched for the presence of cysteine residues (“Total number of cysteines” and “Positions of all cysteines” columns). Proteins with two or more cysteines after the first 30 amino acids of their primary sequence (cells shaded in grey in the “Number of cysteines after position 30” column) are potential substrates of the DSB system for organisms where oxidative protein folding is carried out by DsbA and provided that translocation of the β-lactamase outside the cytoplasm is performed by the Sec system. The first 30 amino acids of each sequence were excluded to avoid considering cysteines that are part of the signal sequence mediating the translocation of these enzymes outside the cytoplasm. Cells shaded in grey in the “Reported in pathogens” column mark β-lactamases that are found in pathogens or organisms capable of causing opportunistic infections. The Ambler class of each enzyme is indicated in the “Ambler class column” and each class (A, B1, B2, B3, C and D) is highlighted with a different color.

**Supplementary Table 2. MIC data used to generate Figure 1B, Figure 1 - figure supplement 2, and Figure 5B.** Cells that are shaded in grey represent strain-antibiotic combinations that were not tested. The aminoglycoside antibiotic gentamicin serves as a control for all strains. For the “Figures 1B and FS2 - MIC values” tab, values are representative of three biological experiments each conducted as a single technical repeat, and for the “Figure 5B - MIC values” tab, values are representative of two biological experiments each conducted as a single technical repeat.

**Supplementary Table 3.** Antibiotic resistance profiles of the clinical isolates tested in this study. The table shows MIC values (µg/mL) for a range of commonly used antibiotics. Values highlighted in pink indicate resistance, as defined by the EUCAST clinical breakpoint guidelines, whilst values highlighted in light blue indicate antibiotics for which there is no EUCAST clinical breakpoint. The remaining values (white cells) indicate sensitivity to the tested antibiotic compound. Strains shaded in yellow are multidrug resistant. The following abbreviations are used: AC, amoxicillin; XM, cefuroxime; TZ, ceftazidime; IP, imipenem; AT, aztreonam; PT, piperacillin/tazobactam; GM, gentamicin; CO, colistin; CI, ciprofloxacin; NF, nitrofurantoin; TR, trimethoprim.

| Strain | AC | XM | TZ | IP | AT | PT | GM | CO | CI | NF | TR |
|---|---|---|---|---|---|---|---|---|---|---|---|
| *E. coli* BM16 (*bla*$_{\text{TEM-1b}}$ *bla*$_{\text{KPC-2}}$) | **>256** | **>256** | **192** | **12** | **>256** | **>64** | 8 | 1 | **>32** | **>512** | **>32** |
| *E. coli* LIL-1 (*bla*$_{\text{TEM-1}}$ *bla*$_{\text{OXA-9}}$ *bla*$_{\text{KPC-2}}$) | **>256** | **>256** | **8** | **3** | **192** | **>64** | 1.5 | 2 | **>32** | 6 | **>32** |
| *E. coli* CNR1790 (*bla*$_{\text{TEM-15}}$ *mcr-1*) | **>256** | **>256** | **32** | 0.5 | **16** | <2 | 1 | **4** | **>32** | 16 | **>32** |
| *E. coli* CNR20140385 (*bla*$_{\text{OXA-48}}$ *mcr-1*) | **>256** | **96** | 1 | 0.25 | 0.38 | **32** | 2 | **4** | **>32** | 8 | **>32** |
| *E. coli* WI2 (*bla*$_{\text{OXA-48}}$ *bla*$_{\text{KPC-28}}$ *mcr-1*) | **>256** | **>256** | **>256** | 1.5 | **32** | **>64** | 1.5 | **4** | 0.016 | 6 | 0.25 |
| *E. coli* 27841 (*bla*$_{\text{CTX-M-55}}$ *mcr-3.2*) | **>256** | **>256** | **16** | 0.19 | **64** | <2 | **32** | **3** | **>32** | 8 | **>32** |
| *E. coli* 1144230 (*bla*$_{\text{CMY-2}}$ *mcr-5*) | **>256** | **96** | **12** | 0.5 | **6** | 8 | 1.5 | **4** | 0.025 | 48 | 1 |
| *K. pneumoniae* ST234 (*bla*$_{\text{SHV-27}}$ *bla*$_{\text{KPC-2}}$) | **>256** | **>256** | **48** | **16** | **128** | **>64** | 0.38 | 2 | 0.047 | **96** | 1 |
| *C. freundii* BM19 (*bla*$_{\text{KPC-2}}$) | **>256** | **>256** | **128** | 4 | **64** | **>64** | **24** | 2 | **>32** | 8 | **>32** |
| *E. cloacae* DUB (*bla*$_{\text{FRI-1}}$) | **>256** | **>256** | **16** | **12** | **>256** | **>64** | 1.5 | **>4** | 0.016 | 48 | 0.75 |
| *P. aeruginosa* PA43417 (*bla*$_{\text{OXA-198}}$) | **>256** | **>256** | 2 | **>32** | 6 | **32** | **16** | 1 | **>32** | **>256** | **>32** |

**Supplementary Table 4.** Bacterial strains used in this study. All listed isolates are clinical strains except for *Escherichia coli* 27841 (ST744), which is an environmental strain. For clinical and environmental isolates, the multi-locus sequence types (ST) are given in parenthesis, where available.

| Name | Description | Source |
|---|---|---|
| ***Escherichia coli*** | | |
| DH5α | $F^-$ *end*A1 *gln*V44 *thi*-1 *rec*A1 *rel*A1 *gyr*A96 *deo*R *nup*G *pur*B20 φ80d*lacZ*ΔM15 Δ(*lac*ZYA-*arg*F)U169 *hsd*R17($r_K^-m_K^+$) $\lambda^-$ | (1) |
| CC118λpir | *ara*D Δ(*ara, leu*) Δ*lac*Z74 *pho*A20 *gal*K *thi*-1 *rsp*E *rpo*B *arg*E *rec*A1 λ*pir* | (2) |
| HB101 | *sup*E44 *hsd*S20 *rec*A13 *ara*-14 *pro*A2 *lac*Y1 *gal*K2 *rps*L20 *xyl*-5 *mtl*-1 | (3) |
| MC1000 | *ara*D139 Δ(*ara, leu*)7697 Δ*lac*X74 *gal*U *gal*K *str*A | (4) |
| MC1000 *dsbA* | *dsbA::aphA*, $Kan^R$ | (5) |
| MC1000 *dsbA att*Tn*7*::P*tac-dsbA* | *dsbA::aphA att*Tn*7*::*dsbA*, $Kan^R$ | This study |
| MG1655 | K-12 $F^-$ $\lambda^-$ *ilv*$G^-$ *rfb*-50 *rph*-1 | (6) |
| MG1655 *dsbA* | *dsbA::aphA*, $Kan^R$ | This study |
| MG1655 *dsbA att*Tn*7*::P*tac-dsbA* | *dsbA::aphA att*Tn*7*::*dsbA*, $Kan^R$ | This study |
| MG1655 *acrA* | *acrA* | This study |
| MG1655 *tolC* | *tolC* | This study |
| MG1655 *degP* | *degP::strAB*, $Str^R$ | This study |
| MG1655 *marR* | *marR::accC*, $Gent^R$ | This study |
| MG1655 *dsbA marR* | *dsbA::aphA marR::accC*, $Kan^R$, $Gent^R$ | This study |
| **Clinical / environmental isolates** | | |
| *Escherichia coli* BM16 | $bla_{TEM\text{-}1b}$ $bla_{KPC\text{-}2}$ | (7) |
| *Escherichia coli* LIL-1 | $bla_{TEM\text{-}1}$ $bla_{OXA\text{-}9}$ $bla_{KPC\text{-}2}$ | (7) |
| *Escherichia coli* CNR1790 | $bla_{TEM\text{-}15}$ *mcr-1* | (8) |
| *Escherichia coli* CNR20140385 | $bla_{OXA\text{-}48}$ *mcr-1* | (8) |
| *Escherichia coli* WI2 (ST1288) | $bla_{OXA\text{-}48}$ $bla_{KPC\text{-}28}$ *mcr-1* | (9) |
| *Escherichia coli* 27841 (ST744) | $bla_{CTX\text{-}M\text{-}55}$ *mcr-3.2* | (10) |
| *Escherichia coli* 1144230 (ST641) | $bla_{CMY\text{-}2}$ *mcr-5* | (11) |
| *Klebsiella pneumoniae* (ST234) | $bla_{SHV\text{-}27}$ $bla_{KPC\text{-}2}$ | (12) |
| *Citrobacter freundii* BM19 | $bla_{KPC\text{-}2}$ | (7) |
| *Enterobacter cloacae* DUB | $bla_{FRI\text{-}1}$ | (13) |
| *Pseudomonas aeruginosa* PA43417 | $bla_{OXA\text{-}198}$ | (14) |
| *Pseudomonas aeruginosa* PA43417 *dsbA1* | *dsbA1* $bla_{OXA\text{-}198}$ | This study |
| *Pseudomonas aeruginosa* PAe191 | $bla_{OXA\text{-}19}$ | (15) |
| *Pseudomonas aeruginosa* PAe191 *dsbA1* | *dsbA1* $bla_{OXA\text{-}19}$ | This study |

**Supplementary Table 5.** Plasmids used in this study.

| Name | Description | Source |
|---|---|---|
| pDM1 | pDM1 vector (GenBank MN128719), p15A *ori*, P*tac* promoter, MCS, $Tet^R$ | Lab stock |
| pDM1-*bla*$_{L2\text{-}1}$ | *bla*$_{L2\text{-}1}$ cloned into pDM1, $Tet^R$ | This study |
| pDM1-*bla*$_{GES\text{-}1}$ | *bla*$_{GES\text{-}1}$ cloned into pDM1, $Tet^R$ | This study |
| pDM1-*bla*$_{GES\text{-}2}$ | *bla*$_{GES\text{-}2}$ cloned into pDM1, $Tet^R$ | This study |
| pDM1-*bla*$_{GES\text{-}11}$ | *bla*$_{GES\text{-}11}$ cloned into pDM1, $Tet^R$ | This study |
| pDM1-*bla*$_{SHV\text{-}27}$ | *bla*$_{SHV\text{-}27}$ cloned into pDM1, $Tet^R$ | This study |
| pDM1-*bla*$_{OXA\text{-}4}$ | *bla*$_{OXA\text{-}4}$ cloned into pDM1, $Tet^R$ | This study |
| pDM1-*bla*$_{OXA\text{-}10}$ | *bla*$_{OXA\text{-}10}$ cloned into pDM1, $Tet^R$ | This study |
| pDM1-*bla*$_{OXA\text{-}198}$ | *bla*$_{OXA\text{-}198}$ cloned into pDM1, $Tet^R$ | This study |
| pDM1-*bla*$_{FRI\text{-}1}$ | *bla*$_{FRI\text{-}1}$ cloned into pDM1, $Tet^R$ | This study |
| pDM1-*bla*$_{L1\text{-}1}$ | *bla*$_{L1\text{-}1}$ cloned into pDM1, $Tet^R$ | This study |
| pDM1-*bla*$_{KPC\text{-}2}$ | *bla*$_{KPC\text{-}2}$ cloned into pDM1, $Tet^R$ | This study |
| pDM1-*bla*$_{KPC\text{-}3}$ | *bla*$_{KPC\text{-}3}$ cloned into pDM1, $Tet^R$ | This study |
| pDM1-*bla*$_{SME\text{-}1}$ | *bla*$_{SME\text{-}1}$ cloned into pDM1, $Tet^R$ | This study |
| pDM1-*mcr-1* | *mcr-1* cloned into pDM1, $Tet^R$ | This study |
| pDM1-*mcr-3* | *mcr-3* cloned into pDM1, $Tet^R$ | This study |
| pDM1-*mcr-3.2* | *mcr-3.2* cloned into pDM1, $Tet^R$ | This study |
| pDM1-*mcr-4* | *mcr-4* cloned into pDM1, $Tet^R$ | This study |
| pDM1-*mcr-5* | *mcr-5* cloned into pDM1, $Tet^R$ | This study |
| pDM1-*mcr-8* | *mcr-8* cloned into pDM1, $Tet^R$ | This study |
| pDM1-*bla*$_{L2\text{-}1}$-StrepII | *bla*$_{L2\text{-}1}$ encoding L2-1 with a C-terminal StrepII tag cloned into pDM1, $Tet^R$ | This study |
| pDM1-*bla*$_{GES\text{-}1}$-StrepII | *bla*$_{GES\text{-}1}$ encoding GES-1 with a C-terminal StrepII tag cloned into pDM1, $Tet^R$ | This study |
| pDM1-StrepII-*bla*$_{OXA\text{-}4}$ | *bla*$_{OXA\text{-}4}$ encoding OXA-4 with an N-terminal StrepII tag cloned into pDM1, $Tet^R$ | This study |
| pDM1-*bla*$_{OXA\text{-}10}$-StrepII | *bla*$_{OXA\text{-}10}$ encoding OXA-10 with a C-terminal StrepII tag cloned into pDM1, $Tet^R$ | This study |
| pDM1-*bla*$_{OXA\text{-}198}$-StrepII | *bla*$_{OXA\text{-}198}$ encoding OXA-198 with a C-terminal StrepII tag cloned into pDM1, $Tet^R$ | This study |
| pDM1-*bla*$_{FRI\text{-}1}$-StrepII | *bla*$_{FRI\text{-}1}$ encoding FRI-1 with a C-terminal StrepII tag cloned into pDM1, $Tet^R$ | This study |
| pDM1-*bla*$_{L1\text{-}1}$-StrepII | *bla*$_{L1\text{-}1}$ encoding L1-1 with a C-terminal StrepII tag cloned into pDM1, $Tet^R$ | This study |
| pDM1-*bla*$_{KPC\text{-}3}$-StrepII | *bla*$_{KPC\text{-}3}$ encoding KPC-3 with a C-terminal StrepII tag cloned into pDM1, $Tet^R$ | This study |
| pDM1-*mcr-1*-StrepII | *bla*$_{MCR\text{-}1}$ encoding MCR-1 with a C-terminal StrepII tag cloned into pDM1, $Tet^R$ | This study |
| pDM1-*mcr-3*-StrepII | *bla*$_{MCR\text{-}3}$ encoding MCR-3 with a C-terminal StrepII tag cloned into pDM1, $Tet^R$ | This study |
| pDM1-*mcr-4*-StrepII | *bla*$_{MCR\text{-}4}$ encoding MCR-4 with a C-terminal StrepII tag cloned into pDM1, $Tet^R$ | This study |
| pDM1-*mcr-5*-StrepII | *bla*$_{MCR\text{-}5}$ encoding MCR-5 with a C-terminal StrepII tag cloned into pDM1, $Tet^R$ | This study |
| pDM1-*mcr-8*-StrepII | *bla*$_{MCR\text{-}8}$ encoding MCR-8 with a C-terminal StrepII tag cloned into pDM1, $Tet^R$ | This study |

| | | |
|---|---|---|
| pGRG25 | Encodes a Tn*7* transposon and *tnsABCD* under the control of P*araB*, thermosensitive pSC101 *ori*, $Amp^R$ | (16) |
| pGRG25-P*tac::dsbA* | P*tac::dsbA* fragment cloned within the Tn*7* of pGRG25; when inserted into the chromosome and the plasmid cured, the strain expresses DsbA upon IPTG induction, $Amp^R$ | This study |
| pSLTS | Thermosensitive pSC101*ori*, P*araB* for λ-Red, P*tetR* for I-SceI, $Amp^R$ | (17) |
| pUltraGFP-GM | Constitutive sfGFP expression from a strong Biofab promoter, p15A *ori*, (template for the *accC* cassette), $Gent^R$ | (18) |
| pKD4 | Conditional oriRγ *ori*, (template for the *aphA* cassette), $Amp^R$ | (19) |
| pCB112 | Inducible *lacZ* expression under the control of the $P_{lac}$ promoter, pBR322 *ori*, $Cam^R$ | (20) |
| pKNG101 | Gene replacement suicide vector, *ori*R6K, *oriT*RK2, *sacB*, (template for the *strAB* cassette), $Str^R$ | (21) |
| pKNG101-*dsbA1* | PCR fragment containing the regions upstream and downstream *P. aeruginosa dsbA1* cloned in pKNG101; when inserted into the chromosome the strain is a merodiploid for *dsbA1* mutant, $Str^R$ | This study |
| pRK600 | Helper plasmid, ColE1 *ori*, *mob*RK2, *tra*RK2, $Cam^R$ | (22) |
| pMA-T *mcr-3* | GeneArt® cloning vector containing *mcr-3*, ColE1 *ori*, (template for *mcr-3*), $Amp^R$ | This study |
| pMK-T *mcr-8* | GeneArt® cloning vector containing *mcr-8*, ColE1 *ori*, (template for *mcr-8*), $Kan^R$ | This study |

**Supplementary Table 6.** Oligonucleotide primers used in this study. The "Brief description" column provides basic information on the primer design (restriction enzyme used for cloning, encoded protein or gene replaced by antibiotic resistance cassette, forward or reverse orientation of the primer (F or R); QC stands for QuickChange primers and SQ stands for sequencing primers).

| Number | Brief description | Sequence (5´-3´) |
|---|---|---|
| P1 | SacI.L2.F | ctggagctcctcgcccgtcgccgatt |
| P2 | XmaI.L2.R | ctgcccgggtcatccgatcaaccggtcggca |
| P3 | SacI.GES.F | ctggagctccgcttcattcacgcac |
| P4 | XmaI.GES.R | ctgcccgggctatttgtccgtgctcaggatg |
| P5 | SacI.SHV.F | ctggagctccgttatattcgcctgtg |
| P6 | XmaI.SHV.R | ctgcccgggttagcgttgccagtgctcga |
| P7 | SacI.OXA-4.F | ctggagctcaaaaacacaatacatataacttcgc |
| P8 | KpnI.OXA-4.R | cagggtaccttataaatttagtgtgtttagaatggtg |
| P9 | SacI.OXA-10.F | ctggagctcaaaacatttgccgcatatgtaattatcgc |
| P10 | KpnI.OXA-10.R | cagggtaccttagccaccaatgatgccctc |
| P11 | NdeI.OXA-198.F | actgcatatgcataaacacatgagtaagctcttc |
| P12 | KpnI.OXA-198.R | ctgggtaccttattcgatgatccccttttgctt |
| P13 | SacI.FRI-1.F | ctggagctctttttttttaaaaaaggtgcaagtac |
| P14 | XmaI.FRI-1.R | ctgcccgggttatttataacttccataaactgcctttatagc |
| P15 | SacI.L1.F | ctggagctccgttctaccctgctcgc |
| P16 | XhoI.L1.R | actgagctctcagcgggccccggccgt |
| P17 | SacI.KPC.F | ctggagctctcactgtatcgccgtc |
| P18 | KpnI.KPC.R | ctgccatggttactgcccgttgacgccca |
| P19 | SacI.SME-1.F | ctggagctctcaaacaaagtaaattttaaaacgg |
| P20 | XmaI.SME-1.R | ctgcccgggttaatcaattgcctgaattgcaatacg |
| P21 | SacI.MCR-1.F | ctggagctcatgcagcatacttctgtgtggtac |
| P22 | XmaI.MCR-1.R | ctgcccgggtcagcggatgaatgcggtgc |
| P23 | NdeI.MCR-3.F | ctgatacatatgccttcccttataaaaataaaaattgttccg |
| P24 | XmaI.MCR-3.R | cagcccgggttattgaacattacgacattgactgaaaatatctag |
| P25 | SacI.MCR-4.F | ctggagctccgtgctgacgagatttaaaaccc |
| P26 | XmaI.MCR-4.R | ctgcccgggttaaccgcggcagcgggcaaaaatatc |
| P27 | SacI.MCR-5.F | ctggagctccggttgtctgcatttatcac |
| P28 | XmaI.MCR-5.R | ctgcccgggtcattgtggttgtccttttctg |
| P29 | SacI.MCR-8.F | ctggagctcttcaagtatcttttatctttcaaact aacc |
| P30 | XmaI.MCR-8.R | ctgcccgggctaaccattcccatctgttttctc |
| P31 | QC.GES5-GES1.F | aaagagccggagatgggcgacaacacacctg |
| P32 | QC.GES5-GES1.R | caggtgtgttgtcgcccatctccggctcttt |
| P33 | QC.KPC2-KPC3.F | ctaacaaggatgacaagtacagcgaggccgtcatc |
| P34 | QC.KPC2-KPC3.R | gatgacggcctcgctgtacttgtcatccttgttag |
| P35 | QC.MCR3-3.2.F | caacgcctttctctttgataaatccaggtgacatcc |
| P36 | QC.MCR3-3.2.R | ggatgtcacctggatttatcaaagagaaaggcgttg |
| P37 | XmaI.StrepII.L2.R | ctgcccgggttatttttcaaattgcggatggctccaagcgctccctccga tcaaccggtcggca |
| P38 | XmaI.StrepII.GES.R | ctgcccgggctatttttcaaattgcggatggctccaagcgctccctttgtc cgtgctcaggatgag |
| P39 | OXA-4.body.F | tcaacagatatctctactgttgca |
| P40 | OXA-4.StrepII.R | tgcaacagtagagatatctgttgatttttcaaattgcggatggctccaagc gctccctgcactggcgctgctgta |

| | | |
|---|---|---|
| P41 | KpnI.StrepII.OXA-10.R | cagggtaccttatttttcaaattgcggatggctccaagcgctcccgccac caatgatgccctcacttg |
| P42 | KpnI.StrepII.OXA-198.R | ctgggtaccttatttttcaaattgcggatggctccaagcgctcccttcgat gatcccctttgcttg |
| P43 | XmaI.StrepII.FRI-1.R | ctgcccgggttatttttcaaattgcggatggctccaagcgctccctttata acttccataaactgcctttatagc |
| P44 | KpnI.StrepII.L1.R | gggggtacctcatttttcaaattgcggatggctccaagcgctcccgcgg gccccggccgtttccttggccaactgc |
| P45 | KpnI.StrepII.KPC.R | ctgccatggttatttttcaaattgcggatggctccaagcgctcccctgcc cgttgacgcccaatc |
| P46 | XmaI.StrepII.MCR-1.R | cagcccgggttatttttcaaattgcggatggctccaagcgctcccgcgg atgaatgcggtgcggt |
| P47 | XmaI.StrepII.MCR-3.R | cagcccgggttatttttcaaattgcggatggctccaagcgctcccttgaa cattacgacattgactgaaaatatctag |
| P48 | XmaI.StrepII.MCR-4.R | ctgcccgggctatttttcaaattgcggatggctccaagcgctcccaccg cggcagcgggcaaaaatatc |
| P49 | XmaI.StrepII.MCR-5.R | ctgcccgggctatttttcaaattgcggatggctccaagcgctcccttgtg gttgtccttttctgca |
| P50 | XmaI.StrepII.MCR-8.R | ctgcccgggctatttttcaaattgcggatggctccaagcgctcccaccat tcccatctgttttctctcttac |
| P51 | NotI.Ptac.EcDsbA.F | ctggcggccgctgacaattaatcatcggctcgtataatgtgtggaattgt gactagtcgaggtccaggacctcggatcgctaagataggatgattgtat gaaaaagatttggctggc |
| P52 | XhoI.EcDsbA.R | ctgctcgagttattttttctcggacagatatttc |
| P53 | EcdsbA::aphA.F | atgaaaaagatttggctggcgctggctggtttagttttagcgtttagcgc gtgtaggctggagctgcttc |
| P54 | EcdsbA::aphA.R | ttattttttctcggacagatatttcactgtatcagcatactgctgaacaagg gaattagccatggtccat |
| P55 | EcacrA::aphA.F | atgaacaaaaacagagggtttacgcctctggcggtcgttctggtgtagg ctggagctgcttc |
| P56 | EcacrA::aphA.R | ttaagacttggactgttcaggctgagcaccgcttgcggcttggggaatt agccatggtccat |
| P57 | EctolC::aphA.F | ttttacagtttgatcgcgctaaatactgcttcaccacaaggaatgcaagtg taggctggagctgcttc |
| P58 | EctolC::aphA.R | tcgtcgtcatcagttacggaaagggttatgatgggaattagccatggtcc |
| P59 | EcdegP::strAB.F | atgaaaaaaaccacattagcactgagtgcactggctctgagtttaggttt ggaactgcacattcgggatatttctc |
| P60 | EcdegP::strAB.R | ttactgcattaacaggtagatggtgctgtcgccgcgctgaatgttgagtg ccaggccggatctagatatctagtatga |
| P61 | EcmarR::accC.F | atggttaatcagaagaaagatcgcctgcttaacgagtatctgtctccgct ggtgaagttcctatactttctagagaataggaacttcaagatcccctg |
| P62 | EcmarR::accC.R | ttacggcaggactttcttaagcaaatactcaagtgttgccacttcgtccgc gaagttcctattctctagaaagtataggaacttcacttactcaatggaattc tagatcg |
| P63 | SQ.dsbA1.Paeruginosa.F | tacctgctcaagcagatgcatg |
| P64 | SQ.dsbA1.Paeruginosa.R | ggtgttcatgtcgcccatca |
| P65 | XbaI.dsbA1.F | ggttcctctagagcctacttcgccagccagaa |
| P66 | dsbA1.body.R | ctacttcttgttacgcatcgttcactc |
| P67 | dsbA1.body.F | atgcgtaacaagaagtaggcaaggtga |
| P68 | BamHI.dsbA1.R | aattaaggatcctcatcactaccaccagcgcg |

**Supplementary Table 7.** Sources of genomic DNA used for amplification of β-lactamase and MCR genes in this study.

| Strain | Gene(s) | Source |
|---|---|---|
| *Stenotrophomonas maltophilia* ATCC 13637 | $bla_{L2\text{-}1}$ $bla_{L1\text{-}1}$ | ATCC |
| *Pseudomonas aeruginosa* GW-1 | $bla_{GES\text{-}2}$ | (23) |
| *Enterobacter cloacae* CHE-2 | $bla_{GES\text{-}5}$ | (24) |
| *Acinetobacter baumannii* K45 | $bla_{GES\text{-}11}$ | (25) |
| *Klebsiella pneumoniae* ST234 | $bla_{SHV\text{-}27}$ $bla_{KPC\text{-}2}$ | (12) |
| *Pseudomonas aeruginosa* SOF1 | $bla_{OXA\text{-}4}$ | (26) |
| *Pseudomonas aeruginosa* PU21 | $bla_{OXA\text{-}10}$ | (27) |
| *Pseudomonas aeruginosa* PA41437 | $bla_{OXA\text{-}198}$ | (14) |
| *Enterobacter cloacae* DUB | $bla_{FRI\text{-}1}$ | (13) |
| *Serratia marcescens* | $bla_{SME\text{-}1}$ | (12) |
| *Escherichia coli* CNR1790 | *mcr-1* | (8) |
| *Shewanella bicestrii* JAB-1 | *mcr-4* | (28) |
| *Escherichia coli* 1144230 | *mcr-5* | (11) |
